# Steller: a high-resolution platform for spatial clonal tracing of mammalian brain development

**DOI:** 10.64898/2026.06.16.732634

**Authors:** Zhuxia Li, Chuwei Wang, Chenyu Guo, Yuanxin Liao, Cheng Chen, Qiaorui Xing, Weike Pei, Guangdun Peng

## Abstract

Resolving both lineage history and spatial position at single-cell resolution is essential for understanding how tissue assemble, yet simultaneously interrogation of both dimensions remains technically challenging. Here, we present Steller (**S**patio-**te**mporal ce**ll** lineage track**er**), an integrated experimental and computational framework that overcomes the sensitivity bottleneck limiting lineage barcode detection in high-resolution spatial transcriptomics (ST). By co-depositing capture probes alongside conventional poly(T) oligonucleotides on high-density spatial arrays, Steller achieves targeted enrichment of lineage barcodes at spatial single cell resolution. Applied to clonally labeled embryonic day 12.5 (E12.5) mouse forebrain development harvested at postnatal day 4 (P4), Steller increased the proportion of barcode-detectable cells and recovered higher clonal diversity compared with poly(T) chip. Leveraging this enhanced sensitivity, we resolved distinct spatial clonal architectures across three forebrain regions: horizontal or radial-to-horizontal transitions in hippocampal pyramidal neurons, dorsoventrally restricted yet multi-nuclei spread in thalamus, and fate-restricted spiny projection neuron lineages. Steller establishes a generalizable strategy for lineage-enhanced spatial profiling compatible with existing high-resolution spatial transcriptomics platforms and adaptable to diverse biologic process, providing a framewrok for investigating lineage-dependent tissue organization.

**Highlights:**

- Steller integrates targeted lineage-barcode enrichment with high-resolution spatial transcriptomics
- Steller substantially improves clonal detection sensitivity at single-cell resolution
- Steller reveals region-specific spatial clonal architectures across the developing mouse forebrain

## Introduction

The functional sophistication of the mammalian brain emerges from the coordinated interplay of neuronal subtype diversification^1,2^ and precise spatial assembly^3,4^. During neurogenesis, cellular identity is orchestrated by the intimate coupling of two fundamental axes: lineage origin—encoding cell-intrinsic developmental potential, and spatial position—conveying extrinsic microenvironmental cues that refine progenitor behavior and terminal differentiation^5^. This interplay manifests across brain regions: whereas E12 cortical progenitors generate radial clonal columns spanning multiple laminae^6^, their counterparts in the hippocampal Cornu Ammonis 1 (CA1) adopt distinct horizontal orientations tailored for functional integration^5^. Similarly, GABAergic fate specification in the basal ganglia is largely predetermined by lineage-intrinsic programs as early as E12.5^7^, whereas the thalamus establish progenitor domains following morphogen gradients^8^ and organize into lineage-dependent nuclei^9^. Disentangling the relative contributions of lineage-intrinsic and extrinsic factors therefore demands methodologies capable of simultaneously interrogating both cellular lineage and spatial position at single-cell resolution.

Recent advances in spatially resolved transcriptomics have transformed our ability to map gene expression directly within intact tissue sections^10^. Platforms ranging from image-based in situ hybridization methods to sequencing-based capture technologies now enable genome-wide expression profiling at resolutions from subcellular to near-single-cell granularity^11^. In parallel, innovations in molecular recording—whereby heritable DNA sequences are introduced into progenitor cells as unique lineage identifiers—have enabled large-scale reconstruction of cellular pedigree relationships^12^. The convergence of these two methodologies—spatial transcriptomics plus clonal barcoding—offers an attractive route toward unified lineage-spatial mapping, as demonstrated by recent studies combining lower-resolution Visium arrays with DNA barcodes^13,14^.

However, translating this combined approach to single-cell spatial resolution encounters a fundamental sensitivity barrier^15^: lineage barcodes introduced via integrating viral vectors constitute small proportion of total polyadenylated mRNA. Conventional spatial capture platforms that rely on poly(T) probes harvest lineage barcodes only incidentally at low efficiency. At higher spatial resolutions—where each capture unit samples a smaller tissue volume containing fewer mRNA molecules—the stochastic probability of barcode capture drops precipitously, framing an undesirable trade-off between spatial resolution and clonal detection rate. Alternative strategies each entail distinct limitations. For instance, imaging-based spatial technologies like SpaceBar^16^ achieve high sensitivity through targeted multiplexed probe hybridization but require predesigned probes for barcode detection, which lack scalability for complex biological programs *in vivo* and unbiased transcriptome co-detection. Computational approaches that infer spatial lineage relationships^17^ from transcriptional similarity elegantly circumvent physical barcode detection but depend on the unvalidated assumption that gene expression state faithfully reflect lineage history—a premise challenged by evidence of rapid transcriptional plasticity during differentiation^6^.

To address this gap, we developed Steller, a framework that combines targeted enrichment of DNA barcodes on high-density spatial arrays with the scalability and whole transcriptomic compatibility of sequencing-based spatial capture. We showed that Steller increased the proportion of barcode-detectable cells and recovered greater clonal diversity, rendering a refined spatially resolved lineage reconstruction across hippocampus, thalamus and striatum at single-cell resolution.

## Results

### Design and validation of the CS3 capture-sequence probe

To address barcode detection limitations inherent to sparse labeling and the sensitivity limitations of high-resolution ST, we designed three distinct CS probes (CS1, CS2 and CS3) complementary to conserved regions of LARRY^18^ lineage barcodes (CloneBC) at the 3’-UTR region of mCherry-CloneBC (Figure 1A and S1A). Probes were linked to conventional poly(T) oligos on spatial bulk read1-ployT beads via hybridization of linker sequences at equimolar input (Figure 1B). Bulk amplicon sequencing of 4T1 cells transfected with the modified LARRY CloneBC library identified CS3 as the optimal probe, exhibiting the highest CloneBC read recovery rate among the three candidates (Figure 1C). To enable joint detection of endogenous transcripts and CloneBC, we functionalized spatial array beads with CS3-modified capture oligos by linking 5’-phosphorylated CS3 probes to the Read1-SpatialBC-UMI-polyT backbone (Figure 1D). Result proves that CS3 substantially outperformed conventional poly(T) in CloneBC read detection efficiency (Figure 1E), indicating that probe enrichment strategy enables spatial transcriptomic chips to gain more comprehensive lineage information at bulk level.

**Figure 1.**
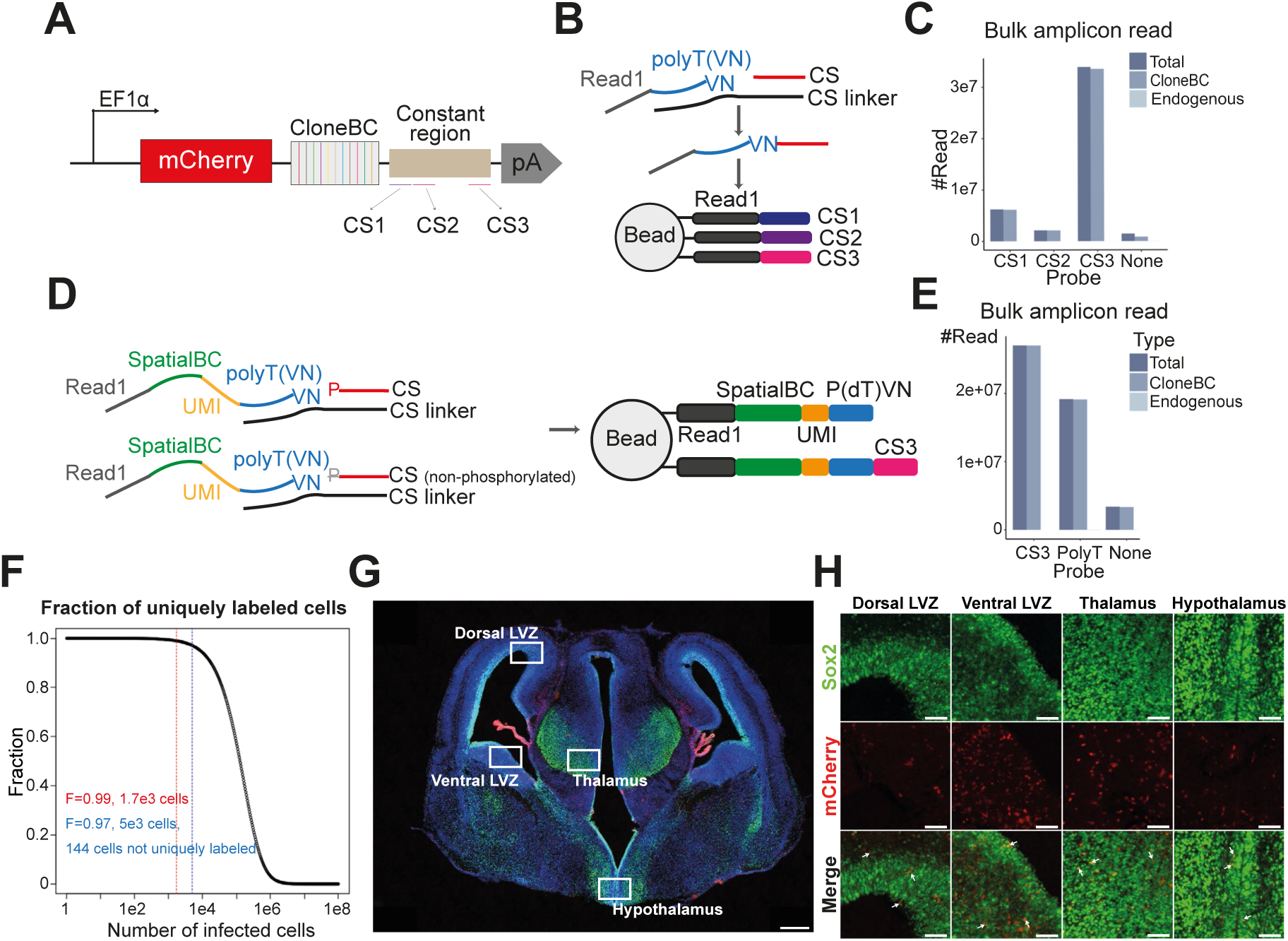
Steller system establishment by integrating CS3-modified ST chip and clonal tracing mouse model. (A-B) Schematic illustration of the binding positions (A) and molecular modifications (B) of the three candidate capture probes designed for CloneBC retrieval. **(C)** Performance benchmarking of the three probe candidates. Bar plots indicate the raw read counts for CloneBC and endogenous mouse transcripts across different CS probe designs. **(D)** Schematic of the partial bead functionalization strategy using a mixture of phosphorylated and non-phosphorylated CS3 probes, highlighting the final structural architecture of the spatial capture oligo. **(E)** Read distribution across poly(T), CS3, and undefined (none) probes derived from bulk amplicon sequencing of CloneBC+ 4T1 tumors. Bar plots illustrating the distribution of read counts (y-axis) across different probe designs (x-axis). For each candidate, total reads are partitioned into those mapping to the exogenous CloneBC and the endogenous mouse transcripts. **(F)** Estimation analysis of clonal labeling uniqueness relative to initial cell numbers. Dashed lines indicate the theoretical 99% uniqueness threshold (red; ∼1,700 cells) and the empirical performance observed in our study (blue; 97% uniqueness at 5,000 cells). **(G)** Representative coronal sections of an E15.5 embryonic brain immunostained for SOX2 (green), 3 days post-CloneBC injection at E12.5 (red: mCherry). **(H)** Magnified view of the boxed region in (G). Arrowheads indicate cells co-expressing mCherry and the SOX2 marker. Scale bars: 300 µm in (G), 50 µm in (H).

Next, we established a lentiviral clonal tracing system for brain development research. The modified CloneBC plasmid library exhibited expected structural architecture, balanced composition and sufficient sequence distance to ensure unambiguous barcode identification (Figure S1B-D). Following *in utero* lateral ventricular microinjecting of low-dose and high-titer CloneBC lentivirus (∼6.12 x 10^6^ TU/μl) at E11.5-E12.5, we achieved sparse labeling with ∼3.14% mCherry^+^ cells (Figure S1E-F). Dual-fluorescence analysis confirmed that labeled cells predominantly contained single copy of CloneBC (Figure S1G). Approximately 97% of labeled cells is estimated to carry unique CloneBC, satisfying the requirement for accurate clonal tracking. The system unbiasedly labeled both SOX2⁺ neural stem cells and SOX2⁻ progenitors of non-neural lineages (Figure 1G–H) and maintained persistent labeling of glial, neuronal, and immune populations through postnatal stages across all forebrain regions examined (Figure S2), validating its suitability for comprehensive developmental lineage analysis.

### Steller captures CloneBC and transcriptome jointly at spatial single cell resolution

To assess Steller’s performance in tissue, we fabricated CS3 probes onto a 2.5 μm ST chip with super-high-density beads for spatial clone analysis at postnal day 4 (P4) following *in utero* E12.5 CloneBC delivery (Figure 2A-B). A customized analytical pipeline was developed to jointly address spatial transcriptome and CloneBC amplicon data (Figure S3A), comprising multi-scale binned matrix construction, cell segmentation and brain region annotation based on transcriptomic data (Figure S2B), as well as quality control, data filtering and lineage related analysis from amplicon data (Figure S2C-D).

**Figure 2.**
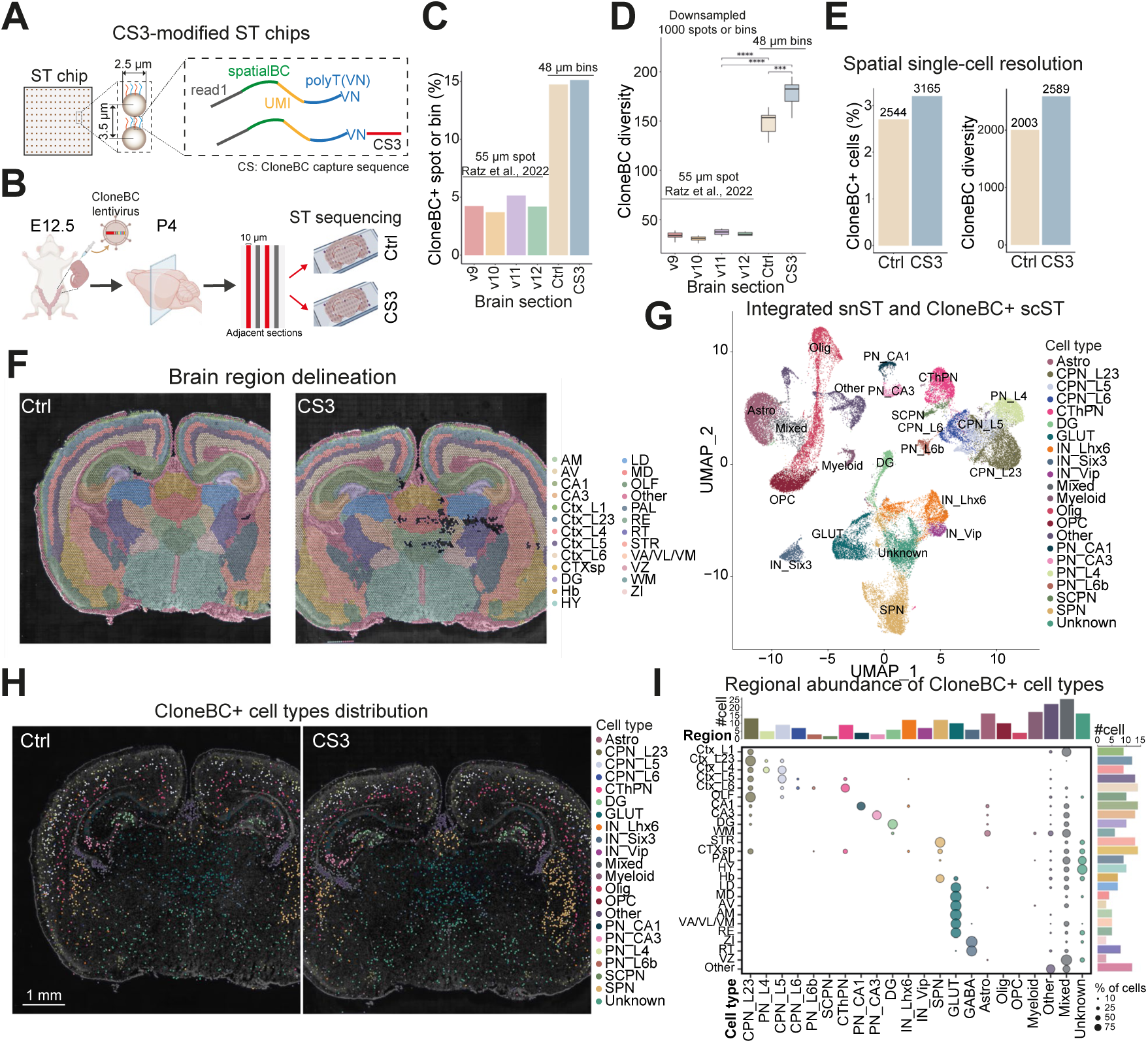
Steller capture CloneBC and transcriptome simultenuously at spatial single cell resolution. **(A)** Schematic of CS3 probe modification on spatial capture oligos. **(B)** Experimental workflow for spatial lineage tracing in the E12.5 mouse brain using control (ctrl) and CS3-modified transcriptome chips. **(C)** Barplot showing the CloneBC+ percentage across Visium spots (55 µm) from Ratz et al., 2022 v9-12 section datasets or bins (level 9, ∼48 µm) of this study. **(D)** Boxplot of total CloneBC diversity at random downsampled 1000 Visium spots or L9 bins with repeats n = 10. Student’s t test was perform between CS3-Ctrl, CS3-v11 and Ctrl-v11 sections. Mean ± SEM; \*\*\**p* < 0.001, \*\*\*\**p* < 0.0001. **(E)** Quantitative comparison of CloneBC capture efficiency at spatial single-cell resolution. Left: Number and percentage of CloneBC+ cells; right: Clonal diversity (number of unique CloneBCs) across ctrl and CS3 chips. **(F)** Level 2 anatomical annotation of the mouse brain visualized via spatial maps. The Cortex (Ctx) is subdivided into distinct laminae, including Ctx_L1, L2/3, L4, L5, L6, and L6b, and olfactory area (OLF). The Hippocampus (HIP) is partitioned into the CA1, CA3, and dentate gyrus (DG). The cerebral nuclei (CNU) is further resolved into the striatum (STR) and pallidum (PAL). The Thalamus (TH) is segmented into discrete nuclei, including the mediodorsal (MD), laterodorsal (LD), anteromedial (AM), anteroventral (AV), reuniens (RE), the ventral anterior/ventrolateral/ventromedial (VA/VL/VM) nucleus, the reticular nucleus (RT), and the zona incerta (ZI). Additional annotated regions include the cortical subplate (CTXsp), hypothalamus (HY), medial habenula (MH), and ventricular zone (VZ). **(G)** UMAP projection of integrated spatial CloneBC+ cells and reference snRNA-seq data, colored by cell-type annotation. **(H)** Spatial maps illustrating the distribution of CloneBC+ cells on ctrl and CS3 chips, overlaid on DAPI-stained nuclei backgrounds. **(I)** Relative cell-type abundance across anatomical brain regions (dot plot), with marginal bar plots indicating total cell counts per cell type (top) and region (right). Scale bars: 1 mm in (H).

At the bin level (48 μm aggregated spatial spots), CS3 chips outperformed concurrently control (ctrl) chips and a published Visium array dataset^14^ in both the proportion of CloneBC-containing spots and CloneBC diversity (Figure 2C-D). Adjusting to single-cell resolution, the proportion of cells with detectable CloneBC (CloneBC^+^ cells) and the diversity of recovered CloneBC (Figure 2E) were both substantially increased in the CS3 chip relative to control, demonstrating that CS3 enhances CloneBC coverage without sacrificing spatial granularity.

We next evaluated whether CS3 probe incorporation compromises endogenous transcriptome fidelity. Total UMI counts and detected gene features showed modest reductions in CS3 chip (Figure SA), but subsequent cluster and cell type analysis remained quite comparable between control and CS3 chips (Figure S4B-C, S5F). Cross-modal concordance between mCherry and CloneBC signals in transcriptome and amplicon data further confirmed data integrity (Figure S4D), indicating that CS3 modification preserve transcriptome-wide profiling quality.

Bin-level transcriptome enabled precise delineation of major forebrain structures (Figure 2F, S5A-C). To overcome sparsity in cell-segmented transcriptome, we integrated ST data of CS3 and control chips with an independently generated spatial single-nucleus RNA-seq (snST) reference dataset, allowing robust cell type classification based on carnonical markers^1,19^ (Figure 2G). Spatial mapping confirmed the distinct distribution of CloneBC^+^ cells across anatomical regions, with relative cell-type abundances consistent with established developmental neuroanatomy (Figure 2H-I, S5D-F). These results further affirming Steller as a sensitive and accurate platform for jointly spatial transcriptome and lineage profiling at single cell resolution.

### Precise lineage relationships identification in tissue context via Steller

Leveraging paired lineage barcodes and spatial transcriptomics, we reconstructed clonal relationships based on the similarity of CloneBC composition within individual cells (Figure S6A-B). Across two sections, we detected clonal information in > 2,000 cells (Figure S6C), with clone size following a Poisson distribution (Figure S6D). Notably, the CS3-modified chip successfully yielded substantially more reconstructed clones and sister-cell pairs than the control chip (Figure 3A). To quantify how lineage origin relates to spatial distribution and cell fate, we computed Euclidean distances between sister-cell pairs in both physical space and transcriptomic space. Sister cells showed greater concordance in both cell fate and spatial location than random cell pairs (Figure 3B), consistent with a model wherein shared progenitors give rise to progeny that occupy overlapping brain regions and adopt similar cell states—a pattern expected under shared exposure to local developmental cues. Specifically, we identified closer clone coupling correlation (Figure 2C) and spatial proximity between transcriptionally similar cell types (Figure S6E). For example, diverse cortical pyramidal neuron (PN) subtypes originated from presumptively the same E12.5 NPC (Figure 3D-E), and clonal sisters tended toward vertical organization, as quantified by the orientation angle between sister-cell pairs (Figure 3F-G), a pattern consistent with radial migration along glial fibers, as reported in traditional sparse fluorescent labeling studies^6^.

**Figure 3.**
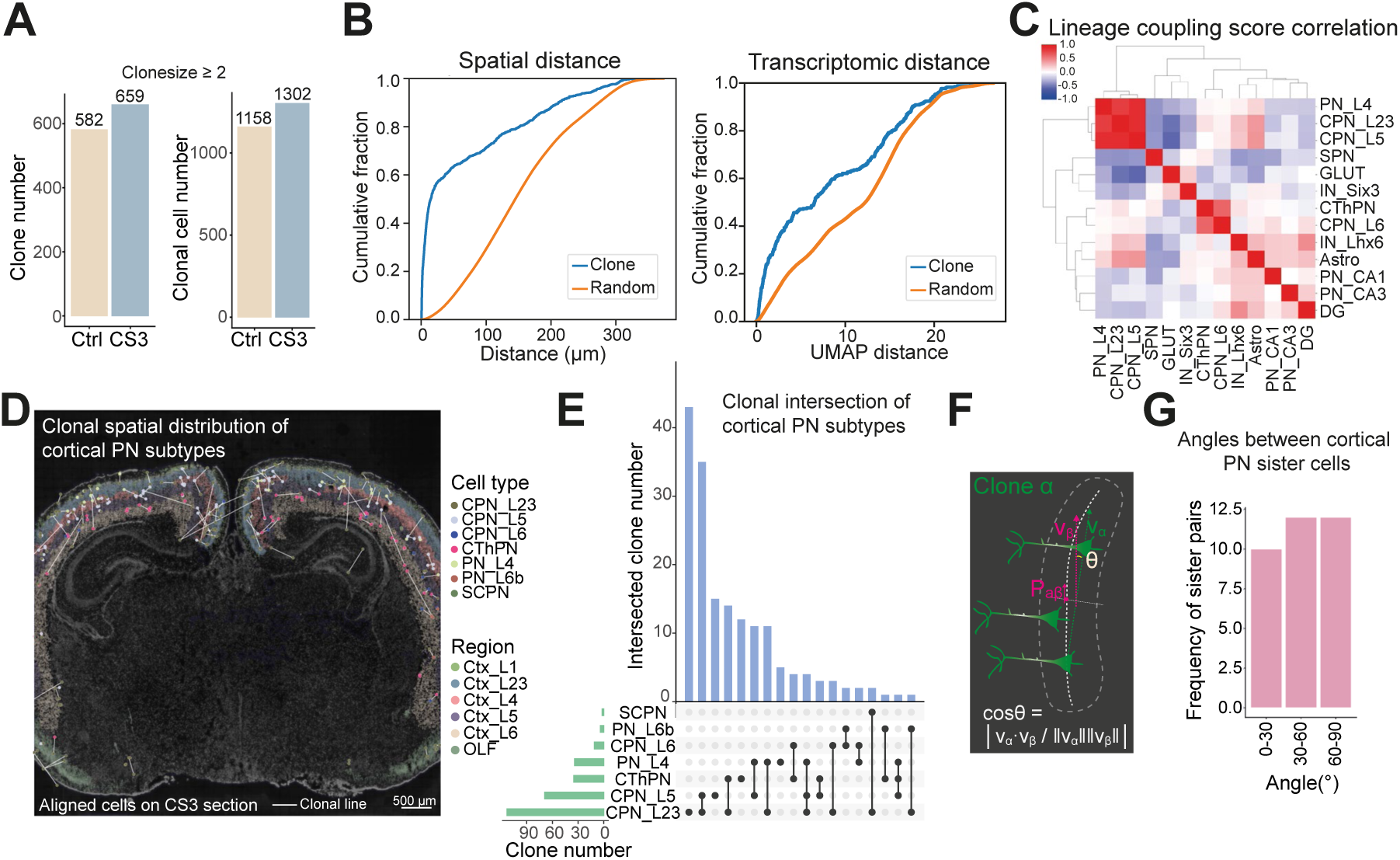
Spatially resolved single-cell lineage with Steller. **(A)** Histgram of clone and clonal cell number detected from ctrl and CS3 sections (for ctrl/CS3, the clone number n = 582/659, clonal cell number n = 1158/1302). **(B)** Cumulative distribution functions (CDFs) comparing the spatial proximity (left) and transcriptomic proximity (Euclidean distance in UMAP space; right) of sister cell pairs versus randomly sampled cell pairs. **(C)** Hierarchical clustering and heatmap of lineage coupling correlation scores among major NPC-derived cell populations. **(D)** Spatial distribution of cortical PN clones across cortical domains. Individual cells are color-coded by cell type; clonal relatives (sister cells) are bridged by white lines. Background shading indicates anatomical brain region annotations. **(E)** UpSet plot illustrating the intersection of shared clones across different PN cell types, highlighting lineage relationships between cortical layers. **(F)** Cartoon representation of calculating the orientation angles (θ) between sister cell pair vector α (Vα) and mid-point nearst tissue horizontal axis vector β (Vβ). **(G)** Orientation analysis of cortical neuronal clones. Histograms quantify the angles between vectors connecting sister cell pairs and the fitted tissue horizontal axis (30° bins). An angle of 90° indicates radial migration, while 0° suggests tangential dispersion along the cortical laminae. Scale bars: 500 µm in (D).

### Diverse spatial lineage-fate modes adopted by distinct brain region development

We next investigated the spatial lineage organization across three anatomically distinct forebrain regions—hippocampus, thalamus and striatum. In the hippocampus, PNs clones in the CA1 region displayed a progressive radial-to-horizontal transition^5^ (Figure 4A-B, S6F-G), whereas clones in dentate gyrus (DG) granule cells and CA3 pyramidal neurons exhibited a predominantly horizontal distribution pattern (Figure 4A-B), which is contrary to the radial arrangement of cortical PNs. These spatial patterns were substantially more discernible in the CS3 group due to its higher abundance of clonal pairs than the control group, particularly in the DG region (Figure 4C). In the thalamus, Steller revealed a distinct pattern: clones were restricted to either dorsal or ventral domains but frequently spread into multiple nuclei within each domain (Figure 4D-E), suggesting early restriction of dorsoventral tangential dispersion at E12.5 but late commitment of progenitors to nucleus-specific fates. In contrast, GABAergic neurons in the reticular nucleus/zona incerta (RT/ZI) showed negligible lineage sharing with glutamatergic (GLUT) neurons in adjacent nuclei (Figure S6H), aligning with the established developmental paradigm about excitatory and inhibitory segregation^20,21^.

**Figure 4.**
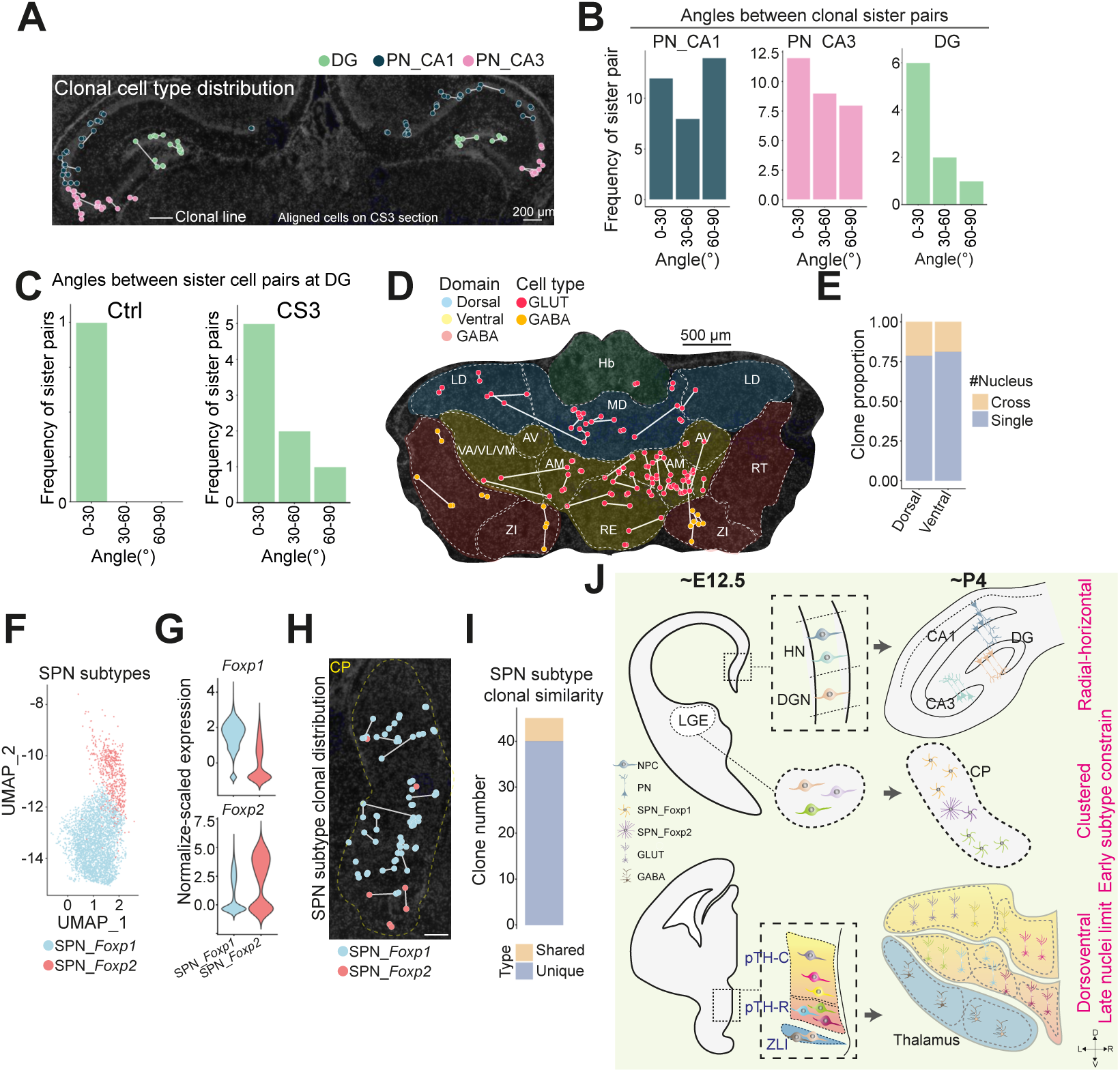
Diverse spatial lineage fate patterns in distinct brain regions. **(A)** Spatial architecture of neuronal clones in the hippocampal formation. Cell types are color-coded, and clonal relatives are connected by white lines. **(B)** Orientation analysis of hippocampal neuronal clones. Histograms show the distribution of angles within each of the three clone subtypes and the fitted tissue horizontal axis (30° bins). **(C)** Histograms of the angles between vectors connecting sister cell pairs and the fitted tissue horizontal axis (30° bins) in ctrl (left) and CS3 (right) sections, respectively. **(D)** Spatial map of thalamic neuronal clones overlaid on anatomical nucleus annotations. Background shading indicates nuclei; dashed lines demarcate dorsal, ventral, and RT/ZI (reticular nucleus/zona incerta) domains. **(E)** Stacked bar plots illustrating the proportion of shared clones across nuclei within the dorsal and ventral thalamic domains. **(F-H)** UMAP projection (F), expression level of *Foxp1* and *Foxp2* (G), and spatial distribution (H) of striatal spiny projection neuron (SPN) subtypes at caudoputamen (CP). Clonal sisters are bridged by white lines over a DAPI-stained background in (H). **(I)** Stacked bar plots showing the number of shared versus non-shared clones between two SPN subtypes. **(J)** Conceptual schematic summarizing spatial clonal lineage patterns across the hippocampus subregions, thalamus, and caudoputamen. Clones are color-coded.pTH-C: thalamic caudal domain; pTH-R: thalamic rostral domain; ZLI: zona limitans intrathalamica. Scale bars: 200 µm in (A), 500 µm in (D) and 200 µm in (H).

Finally, within the caudoputamen (CP) of striatum, we identified two spiny projection neuron (SPN) subtypes, which can be molecularly distinguished as *Foxp1*^+^ and *Foxp2*^+^ populations (Figure 4F-G). Spatial mapping showed that although SPN clones were locally clustered (Figure 4H, S6E), they exhibited notable fate restriction patterns: sister cells overwhelmingly belonged to the same SPN subtype (*Foxp1*^+^ or *Foxp2*^+^), with very few clones containing a mixture of both (Figure 4I). This observation was also validated by reanalyzing a previous published single cell lineage tracing dataset (Figure S6I-J). It indicates that lineage history is tightly coupled to the acquisition of specific molecular identities during striatal development, which may be functionally linked to the establishment of specialized dopaminergic neurotransmission circuits^22,23^.

Collectively, these region-specific lineage patterns—horizontal orientation in the hippocampus, domain-confined propagation in the thalamus, and subtype-restricted clustering in striatum—highlight how conserved developmental machinery, such as radial glia guidance and morphorgen cues, is spatially deployed to build functionally distinct brain structures (Figure 4J).

## Discussion

We have presented Steller, an integrated framework achieving sensitive detection of lineage barcodes in high-resolution spatial transcriptomics through capture-sequence probe-mediated targeted enrichment. By restoring barcode detection rates to levels supporting robust clonal reconstruction at single-cell spatial resolution, Steller bridges a methodological gap between existing low-resolution spatial-barcode approaches^13,14^ and non-spatial single-cell lineage tracing methods^18^, enabling—to our knowledge, for the first time—systematic interrogation of spatial clonal architecture across multiple anatomical regions within a single tissue specimen. Its reliance on standard reverse-transcription-based capture chemistry ensures compatibility with existing commercial and academic spatial transcriptomics platforms; its probe-based enrichment strategy is agnostic to barcode library composition (requiring only knowledge of conserved flanking sequences); and its retention of full transcriptome co-detection enables joint analysis of lineage identity and cell-state markers within the same cells. Steller increased the proportion of CloneBC-detectable cells and recovered substantially greater clonal information compared with conventional poly(T)-based capture—performance gains that proved essential for resolving fine-grained spatial clonal architectures.

Application to E12.5 mouse forebrain development unmasked divergent lineage-spatial organization principles at newborn stage. In the hippocampus, subregional divergence was pronounced: CA1 pyramidal neuron clones transitioned from radial to horizontal distributions, suggesting extended tangential dispersion following initial radial migration, whereas CA3 pyramidal neurons and DG granule cells exhibited predominantly horizontal orientations. These distinct geometries likely underpin the specific lineage-dependent structural and functional organization of hippocampal subcircuits ^5^. In the thalamus, clones were strictly dorsoventrally partitioned at E12.5 yet subsequently dispersed across multiple nuclei, indicating early spatial restriction followed by late nucleus-specific fate commitment. This dorsoventral patterning potentially inherited the early embryonic signaling cues and directs subsequent cortical connectivity and relay functions^8,20^. The striatum displayed the moust striking logic: SPN clones exhibited tight spatial clustering coupled with near-absolute molecular fate restriction. Sister cells almost invariably adopted the same *Foxp1*⁺ or *Foxp2*⁺ identity, indicating early subtype specification. This early embryonic divergence likely predetermines the functional segregation of direct and indirect pathways within the basal ganglia circuit^22,23^.

These findings raise the possibility that the timing of lineage-dependent progenitor specification is regionally variable. We hypothesize that in the striatum, subtype identity is specified prior the final mitotic division, effectively assigning fate within specific local environment. Conversely, in the hippocampus and thalamus, greater post-mitotic dispersion is permitted, allowing for a more fluid integration of lineage into spatial architectures. This hypothesis predicts that experimental manipulation of progenitor specification timing would differentially alter spatial clonal architecture across regions—a prediction that can now be rigorously tested using Steller in combination with temporal control of progenitor manipulation (e.g., *in utero* electroporation at different embryonic time points).

Collectively, Steller provides a generalizable strategy compatible with existing high-resolution spatial transcriptomics platforms and extensible to diverse biological processes including tumor evolution, tissue regeneration, aging and stem cell dynamics. As tissue section throughput increases and multi-temporal sampling is integrated, Steller is poised to deliver exponential gains in lineage information, facilitating the transition toward a comprehensive 3D, and ultimately 4D (spatiotemporal), single-cell lineage atlas of mammalian fate mapping.

### Limitations

Several limitations should be considered when interpreting these findings. First, the analysis derives from a limited number of biological replicates and tissue sections, which constrains statistical power for detecting subtle regional differences. Second, lineage relationships were static and reconstructed from two-dimensional sections, which capture only a subset of three-dimensional clonal architectures and may overlook dispersal patterns orthogonal to the sectioning plane. Third, non-neuronal populations—including glia, immune cells, and other lineages—were underrepresented in the dataset, limiting our ability to assess lineage-spatial relationships across the full cellular diversity of the developing forebrain. Fourth, the single postnatal time point precludes direct reconstruction of dynamic migratory trajectories and temporal fate transitions. Finally, the molecular and cellular mechanisms underlying clonal spatial organization and fate specification remain to be elucidated. Future extensions incorporating serial-section three-dimensional spatial lineage reconstruction, time-course sampling across developmental stages, and correlation with neural activity-dependent refinement processes will be essential for building a mechanistic understanding of how clonal origin instructs brain wiring and function.

## STAR Methods

### Generation of CloneBC plasmid library and lentivirus

#### CloneBC plasmid library generation

LARRY barcode library was generated following Weinreb et al., 2020^18^ with modification. To achieve efficient fluorescent labeling for in vivo ventricular infection, the EF1a-mCherry expression cassette was inserted in the forward orientation between the Long Terminal Repeat (LTR) sequences of the lentiviral backbone. This orientation was optimized to enhance viral transcript packaging efficiency. To generate the LARRY barcode library, equimolar amounts of forward and reverse single-stranded oligos (10 μM each, LARRY-F and LARRY-R, see Supplementary Table 1) were annealed in 1× NEBuffer 4 (NEB, B7004S) using a temperature gradient (95°C for 5 min, followed by-0.1°C/s ramps to 65°C for 5min and then followed by-0.1°C/s ramps to 4°C). The annealed products were extended using Klenow Fragment (NEB, M0212L) at 37°C for 2 h, followed by enzyme inactivation at 70°C for 20min and a hold at 4°C. After purification (Monarch PCR & DNA Cleanup Kit, NEB, T1030S), approximately 2 μg of the extended products was double-digested with BamHI-HF (NEB, R3136S) and XbaI (NEB, R0145S) at 37°C for 12 h. The digestion was terminated by 10% SDS and the digested oligos were purified and normalized to a concentration of 1 ng/μL. In parallel, the lentiviral backbone (∼8.5 μg) was linearized via BamHI-HF and XbaI digestion for 4 h, followed by dephosphorylation with rSAP (NEB, M0371S) for an additional 4h incubation at 37°C. The linearized vector was purified and ligated with the digested LARRY oligos (200 ng vector: 4.5 ng insert) using T4 DNA Ligase (NEB, M0202S) at 16°C for 16 h. The ligation product was purified and transformed into MegaX DH10B T1R Electrocomp cells (Invitrogen, C640003) via electroporation (2000 V, 200 Ω, 25 μF, BioRad, GenePulserXcell). After recovery in SOC medium, approximately 10 μL of the suspension was serially diluted several thousand-fold and plated to calculate the library complexity, while the remaining bacterial was plated on LB agar plates containing 80 μg/mL ampicillin. All colonies were scrapped, pooled and subjected to plasmid extraction using a midiprep kit (Invitrogen, K210014), yielding a final plasmid library of 100–350 μg.

#### Lentiviral production and concentration

Lentiviral vectors were packaged using a second-generation three-plasmid system in HEK293T cells. Briefly, 293T cells were cultured in DMEM supplemented with 10% FBS on poly-D-lysine (PDL, Gibco, A3890401)-coated 10-cm dishes. Upon reaching 90–95% confluence, cells in each dish were co-transfected with 4 μg psPAX2, 2 μg pMD2.G, and 4 μg CloneBC plasmid library using 40 μg polyethylenimine (PEI, Yeasen, 40816ES02) in serum-free Opti-MEM medium (Gibco, 31985070). The medium was replaced with Opti-MEM containing 10% fetal bovine serum (FBS) at 12h post-transfection. Viral supernatants were harvested every 12h with centrifugation of 1,500 x g for 5 min. The pooled supernatant was concentrated by ultracentrifugation at 50,000 x g for 2h at 4°C (JA30.50 rotor, Beckman Coulter, Avanti J-30i). After aspirating the supernatant, the viral pellets were resuspended in a specialized resuspension buffer (20 mM Tris-HCl, 250 mM NaCl, 10 mM MgCl, and 5% sorbitol) and stored in aliquots at-80°C.

#### Lentiviral titration

Viral titers were determined by infecting 293T cells. Cells were seeded at 4 x 10^5 cells per well in 6-well plates and transduced with a serial dilution of the concentrated virus in the presence of polybrene (8 μg/mL, OriLeaf, S24797). After 72 h of incubation, cells were harvested and the percentage of mCherry-positive cells was quantified by flow cytometry (Beckman Coulter, CytoFlex). The infectious titer (expressed as transducing units per mL, TU/mL) was calculated using the following formula:

Titer = (F x N x D) x 1000, where F is the fraction of mCherry^+^ cells, N is the cell number at the time of transduction, and D is the dilution factor.

### CloneBC whitelist determination

To characterize the CloneBC diversity within the plasmid library and facilitate downstream data filtering, a representative 50∼80 ng aliquot of the plasmid library was subjected to enrichment PCR for verification of barcode whitelist. The reaction was performed in a 50 μL system containing 25 μL of 2× KAPA HiFi Mix (Roche, KK2602) and 1.5 μM each of indexed forward and reverse primers (F-primer and R-primer, 10 μM stock). The amplification involved an initial denaturation at 95°C for 3 min, followed by 8 cycles of 98°C for 20 s, 60°C for 15 s, and 72°C for 15 s, with a final extension at 72°C for 5 min. The resulting PCR products were purified using the Monarch PCR & DNA Cleanup Kit with a 7:1 binding buffer ratio and quantified via Qubit (Thermo Fisher Scientific, Q33226) fluorometric assay. Sequencing libraries were prepared using the VAHTS Universal DNA Library Prep Kit for Illumina V3 (Vazyme, ND607-01) according to the manufacturer’s instructions. High-throughput sequencing was executed on the Illumina NovaSeq 6000 platform in PE150 mode to achieve sufficient coverage of the barcode library. For primer sequences and indices, see Supplementary Table 1.

### Mice

7∼12 weeks old adult ICR mice were purchased from Zhuhai BesTest BioTech Co,.Ltd. The day of vaginal plug detection was designated as E0.5, and the day of birth was designated as postnatal day 0 (P0). All experiments were performed under the approval of the Animal Care and Use committee of Guangzhou Institutes of Biomedicine and Health, Chinese Academy of Sciences (Institutional Animal Welfare Assurance Number A5748-01).

### CloneBC^+^ 4T1 tumor RNA preparation

The 4T1 mouse breast cancer cell line was provided by Weike Pei lab from Westlake University. 4T1 cells were firstly transduced with high-titer CloneBC lentivirus to achieve nearly 100% positivity. For implantation, 1 x 10^6 cells were resuspended in a 1:1 mixture of PBS and Matrigel (100 μL total volume, Corning, 354230) and kept on ice prior to injection. The cell suspension was injected into the right fourth mammary fat pad of anesthetized mice. To ensure high-efficiency engraftment and minimize cell leakage, the injection site was secured with forceps for 1 min after needle withdrawal. Tumors were harvested 14 days post-injection, ensuring the maximum diameter remained below 1 cm in compliance with institutional animal welfare protocols. Excised tumor tissues were weighed, measured, and immediately embedded in OCT compound for storage at-80°C. For modified CS probe test, 5∼10 50 μm cryosections were pooled. Total RNA was extracted from these sections with RNA concentration quantified via Qubit fluorometry and RNA integrity (RIN value) assessed using an Agilent Bioanalyzer.

### *In utero* microinjection of CloneBC lentivirus

Glass capillaries (Sutter, B100-50-10) were prepared using flaming/brown micropipette puller (Sutter Instrument, Model P-1000) (Heater: 671, Pull: 190, Velocity: 120, Delay: 250, Pressure: 500). The pulled capillaries were manually trimmed and beveled into needles with an outer diameter of 40~45 μm. Needle tips was sharped using a micro-grinder (KeDouBC, KD-MG3) via sequential grinding at descending angles (30°, 25°, and 20°) for 2~5 min per stage. Before use, needles were cleaned by ethanol aspiration and calibrated using a micro-injection pump (Eppendorf, FemtoJet 4x). The injection pressure required to deliver a 0.6 μL volume of phosphate-buffered saline (PBS) was recorded for each capillary to ensure consistent intraventricular delivery. Timed-pregnant mice were anesthetized with isoflurane until the loss of pedal reflex. Following abdominal depilation and skin disinfection, a 1∼2 cm midline laparotomy was performed to expose the uterine horns. To ensure high maternal and fetal survival rates, the entire surgical duration for each mouse was restricted to under 30 min. CloneBC lentiviral inoculum was mixed with 1/10 volume of 0.1% Fast Green dye for injection visualization. The injection needle was advanced through the uterine wall and amnion, and successful placement into the fetal lateral ventricle was confirmed by the rapid diffusion of the dye within the ventricular space. After injection, the abdominal muscle layer was sutured and the skin incision was closed with surgical staples. Mice were maintained at 37°C on a heating pad during recovery and administered subcutaneous analgesia for three consecutive days. Postoperative recovery and health status of the dams were monitored daily until the birth of viable experimental pups.

### Immunofluorescence staining of mouse brain sections

After dissection, the brain tissue was fixed in 4% paraformaldehyde (PFA) overnight at 4℃, followed by cryoprotection in 30% sucrose dissolved in 1 x PBS at 4°C until the tissue sank to the bottom of the tube. Tissues were embedded in OCT compound within cryo-mold on dry ice and stored at −80°C. Frozen sections were cut at 10∼20 µm thickness, mounted onto slides, and air-dried for 15∼20 minutes at room temperature. For frozen samples, sections were post-fixed with 50 µL of ice-cold 4% PFA for 5 minutes at room temperature, followed by three washes in PBS-Triton Wash Buffer for 5 minutes per wash. Sections were incubated with 30 µL primary antibodies diluted in blocking buffer and covered with parafilm in a humidified chamber overnight at 4°C. List of primary antibodies are as follows: SOX2 (rabbit, 1:500, CST, 23064s), mCherry (rabbit, 1:500, Abcam, ab167453) PAX6 (mouse, 1:500, Abcam, ab78545), NEUN (rabbit, 1:500, Millipore, ABN78), GFAP (rabbit, 1:200, Abcam, ab278054), OLIG2 (rabbit, 1:500, Sigma, AB9610) or IBA1 (chicken, 1:500, Synaptic Systems, 234009). Following primary antibody incubation, sections were washed 3 times for 15 minutes each in 1X PBS. Sections were then incubated with 50uL fluorophore-conjugated secondary antibodies diluted in blocking buffer against the respective species (donkey-anti-rabbit Alexa Fluor 488, 1:500, Abcam, ab150073; donkey-anti-mouse Alexa Fluor 647, Abcam, ab150107; Alexa Fluor 488 AffiniPure Donkey Anti-Chicken, Jackson, 703-545-155), under parafilm in a humidified chamber for 2.5 h at room temperature, protected from light. After secondary antibody incubation, sections were washed 3 times for 5 minutes each in 1× PBS. Nuclei were counterstained with 30 µL of 1 μg/mL DAPI for 2 minutes at room temperature, followed by a single rinse in 1× PBS for 10∼15 minutes. Sections were mounted with an anti-fade mounting media, avoiding bubbles. Fluorescence images were acquired using microscope (TissueGnostics, Tissue FAXS Plus ST).

### CloneBC uniquely labeling estimation

To estimate the initial number of progenitor cells labeled at the time of microinjection *Nt*0, we employed a mathematical model^8,24^ based on the cell population at a subsequent time point *Nt*1. Given the time interval Δ*t* between injection and analysis, and the daily cell division frequency *f*, the initial transduced population was determined by:

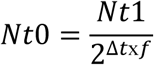

Based on flow cytometric analysis of E15.5 embryonic mouse brains, we observed an average mCherry positive rate of 3.14% among a total of 2.927 x 10^6 cells, yielding *Nt*1 = 91,900. Using a recorded division frequency^25^ *f* = 1.4 and Δ*t* = 3 for neural stem cells between E12.5 and E15.5, the initial labeled population *Nt*1 was estimated to be approximately 5,000 cells.

To account for the non-uniform distribution (Gini coefficient) of our plasmid library, we calculated the expected number of non-uniquely labeled cells *E*(*X*) using the following probabilistic model:

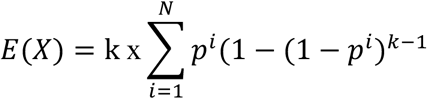

where *p^i^* represents the frequency of a specific CloneBC *i* = 1…*N* within the library, and 9 is the number of initially infected cells. Given our typical injection of *N* = 6.12 x 10^6^ barcodes and an estimated *k* = 5000 cells, the model predicted. *E*(*X*) = 144 non-uniquely labeled cells. This corresponds to a 97% unique labeling efficiency, confirming the high resolution of our lineage tracing system.

### CloneBC copy number estimation

mScarlet and mBaoJin fluorescent reporters (donated by Zhifei Fu’s laboratory from Fujian Medical University) were subcloned into the lentiviral backbone via restriction digestion and ligation. Lentiviral particles were packaged and concentrated according to the aforementioned protocols, achieving a final titer in the range of 10^9 infectious units/mL. The two fluorescent viruses were mixed at an equimolar ratio and microinjected (total volume matching experimental conditions) into the lateral ventricles of E11.5 embryos. At 3∼4 days post-injection (E14.5∼E15.5), embryonic brains were harvested and enzymatically dissociated using papain (Worthington, LK003153). The resulting single-cell suspensions were analyzed via flow cytometry to quantify the proportions of single-positive (mScarlet^+^ or mBaoJin^+^) and double-positive (mScarlet^+^/mBaoJin^+^) cell populations.

### Modification of CS probes on BMKMAN S3000 chip (BMKMANU company)

For bulk-level bead functionalization, a mixture of CS1, CS2, and CS3 probes was conjugated to read1-polyT beads at equimolar ratios. This was achieved using a specialized linker containing sequences that are fully and partially reverse-complementary to the probes and the read1-polyT handle, respectively. Detailed sequences for all probes and linkers are provided in the Supplementary Table 1.

On the spatial chip, a mixture consisting of 60% phosphorylated and 40% non-phosphorylated CS3 probes was linked to read1-SpatialBC-UMI-polyT oligos immobilized on the beads. To ensure the specific retention of functionalized templates, the non-phosphorylated probes were removed via a potassium hydroxide (KOH) washing step. This optimization protocol resulted in approximately 50% of the total capture probes being successfully functionalized with the CS3 sequence.

### Tissue preparation of spatial transcriptomics

Postnatal day 4 (P4) mice, previously subjected to in utero microinjection at E11.5∼E12.5, were sacrificed, and brain tissues were rapidly frozen. Sequential coronal sections (10 µm thickness) were obtained starting from the bregma zero coordinate (BZC)-1 mm. Two representative sections, separated by a single-section interval, were selected for BMKMANU spatial RNA-seq. In parallel, to provide higher-order anatomical and confident transcriptomic context at real single cell level, the left hemisphere of a P13 mouse brain was sectioned at 20 µm thickness within a comparable anteroposterior (A-P) interval and subjected to SeekSpace sequencing.

### Spatial transcriptomics and amplicon library generation and sequencing

For spatial RNAseq transcriptomic library, the 2.5 μm BMKS3000 platform was utilized and spatial transcriptome analysis was done following the manufacturer’s instructions (the BMKMANU Tissue Optimization Reagent Kits User Guide V2.1, BMKMANU Gene Expression kits user guide V3.4 and BMKMANU S Library Construction Kit User Guide V3.1). Libraries were sequenced on the GeneMind SURFseq5000 platform with PE150 mode.

For spatial snRNAseq transcriptomic library, SeekSpace platform (SeekGene company) was applied according to the manufacturer’s instructions (SeekSpace Single Cell Spatial Transcriptome-seq Kit (K02501-08)). The single-cell RNA-Seq library and spatial barcode library were then sequenced on the Illumina NovaSeq X Plus with PE150 read length. The CloneBC library was retrieved from spatial RNA and enriched via a half-nested PCR strategy using KAPA HiFi HotStart ReadyMix (Roche, KK2602). The primary amplification (50 μL reaction) used 50 ng of cDNA template with 1.5 μL of 10 μM F-2410 forward primer and 1.5 μL of 10 μM TruSeq-Read1 reverse primer. PCR program consisted of an initial denaturation at 95°C for 3 min, followed by 6–10 cycles of 98°C for 20 s, 62°C for 15 s, and 72°C for 35 s, with a final extension at 72°C for 5 min. The first PCR products were purified using left side SPRI selection (0.8x) with VAHTS DNA Clean Beads (Vazyme, N411-01) and eluted in 20 μL of nuclease-free water. The entire purified volume served as the template for the second PCR, utilizing TruSeq-R2_F forward and TruSeq-Read1 reverse primers. This reaction followed a similar amplification procedure but was extended for 8 cycles with a 30 s elongation step at 72°C. Secondary products were again purified using 0.8x left side SPRI selection and quantified via Qubit. Final library indices and Illumina adapters were incorporated through a terminal 8-cycle PCR using 50 ng of the half-nested product, Illumina P5 and P7 primer. Cleaned library was analyzed by Qubit and Bio-Fragment Analyzer (Bioptic, Qsep100). Libraries were sequenced on the SURFseq5000 with PE150 read length. For primer sequences and indices, see Supplementary Table 1.

### CS probe selection and read type identification

Total RNA from 4T1 tumors was hybridized with beads modified with CS1/2/3 probes as previously described. cDNA was synthesized and amplified using a Template Switching Oligo (TSO) and TruSeq-Read1 primers, followed by amplicon library preparation and high-throughput sequencing as described later. Raw reads were quality-filtered (FastQC, > Q25). To identify the capture probe type, the first 26 bp of Read 1 were aligned against CS1, CS2, and CS3 sequences using PairwiseAligner score with parameters (mode = “local”, match_score = 1, mismatch_score =-0.5, open_gap_score =-0.5, extend_gap_score =-0.5) from BioPython package (v1.85). Probes were assigned based on the score exceed 16.5, 20.5 or 18.5 for CS1, CS2 and CS3, respectively. Reads failing to meet this threshold were categorized as other.

### Spatial transcriptomics data analysis

#### Cell segmentation

To achieve single-cell resolution from the spatial transcriptomic data, nuclear staining images were utilized for the precise segmentation of individual spatial cells. This process was executed using the Cellpose algorithm, which was integrated into the BSTMatrix pipeline (v2.4.e). The segmentation utilized a customized parameter set to optimize detection in brain tissue (Diameter: 28, Cell Edge: 1, Minimum Detection Value: 50, Flow Threshold: 0.2). Following segmentation, the aligned spatial transcriptomic signals were partitioned based on the identified cellular boundaries. This transformation converted the spot-based data into a spatial single-cell transcriptomic matrix, facilitating downstream analyses that require cellular-level granularity, such as cell-type-specific lineage tracing and high-resolution spatial clustering.

#### Brain region identification

Clustering was performed based on gene expression matrix at level 9 (about 48 μm resolution), clustering was performed. According to previous canonical and known markers^1,26–30^ and public early postnatal development mouse brain atlas at P04 age^31^ (https://kimlab.io/brain-map/epDevAtlas/), brain region was annotated.

#### Cell type annotation

The CloneBC^+^ spatial cellular transcriptomic matrix and the spatial single-nucleus expression matrix were integrated using Canonical Correlation Analysis (CCA). This cross-platform integration allowed for the alignment of transcriptomic states across the experimental (P4) and reference (P13) datasets. Following integration, we performed unsupervised clustering on the joint dataset. Clusters were manually annotated based on two primary criteria: 1) Differential expression analysis was used to identify cluster-specific marker genes and functional signatures. 2) The anatomical distribution of each cluster was mapped back onto the original tissue coordinates to ensure consistency with known brain architecture.

### Spatial amplicon data analysis

#### Preprocessing

The raw fastq files were first processed through BSTMatrix pipeline (steps 1∼5) to generate the initial Level 1 (L1) gene expression matrix and identify spatial spot. Using the previously generated cells.npy (nuclear segmentation) and the L1 spots, the mapping relation file of L1 spot and individual segmented cells was generated.

CloneBC were retrieved from the sequencing reads by the following steps: 1) Reads were first aligned to the mCherry in the BAM files. 2) To minimize technical artifacts, UMIs with a total read count of one were discarded. 3) CloneBC sequences were called using a specific regular pattern:

[ATCG]{4}CT[ATCG]{4}AC[ATCG]{4}TC[ATCG]{4}GT[ATCG]{4}TG[ATCG]{4}CA[ ATCG]{4}

Extracted CloneBC-UMI pairs were mapped back to individual cells based on the L1 spot-to-cell spatial relationship.

Besides, in order to generate a high-confidence spatial clone map, we applied rigorous filtering at the cell and barcode levels: 1) Cells with total reads < 10 and specific cell-CloneBC pairs with reads < 5 were removed. 2) CloneBC sequences were collapsed using Starcode (default distance) to merge related sequences likely originating from PCR or sequencing errors. 3) To eliminate high-frequency technical noise or ambient barcode contamination, we implemented a two-tier “confidence” metric: a. Confident CloneBC: defined as a CloneBC accounting for > 10% of the total CloneBC UMIs within a specific cell (CloneBC_umi_percent_percell > 0.1). b. Confident cell: defined as a cell where > 50 of its UMIs belong to “confident” CloneBCs (conf_CloneBC_percent > 0.5). Only CloneBCs and cells meeting these confidence thresholds were retained for downstream analysis.

Following the merging of CloneBC-cell matrices from the two adjacent tissue sections, the dataset was subjected to outlier removal based on standard deviation (s.d.) thresholds: 1) To eliminate potential doublets or cells with non-specific barcode accumulation, cells exhibiting an abnormally high CloneBC UMI count (nCount_CloneBC > 1.5*s.d. above the mean) were excluded. 2) To remove ubiquitous barcodes, CloneBCs that appeared with excessive frequency across the cell population (nCell > 2*s.d. above the mean) were discarded.

#### Clone calling

The cleaned and merged matrix is used to define clones. Clonal identity was determined by assessing the similarity of barcode combinations between cells. The Jaccard similarity coefficient was calculated for every pair of CloneBC-expressing cells to quantify the overlap of their respective barcode sets. A stringent Jaccard score cutoff of 0.7 was applied; cell pairs exceeding this threshold were considered significantly related and connected within the similarity network. Cells were modeled as nodes in an undirected graph, with edges representing connections that passed the Jaccard threshold. Clones were defined as the connected components of the graph, specifically consisting of clusters of two or more related cells.

### Image data analysis

Nuclear staining images from two adjacent sections were manually aligned and subjected to non-rigid registration. The resulting registration coordinates were applied to the segmented cell centroids, ensuring precise spatial alignment of cellular identities across sections. Finally, the registered cells from the adjacent layer were overlaid onto the CS3 section for integrated visualization of cell type distribution and clone relations.

### Lineage coupling correlation score

To evaluate lineage proximity between cell types, we calculated coupling Z-scores following the framework described by Bandler et al^7^. Cell types with fewer than 20 clonal cells were excluded. Briefly, a clone was defined as “shared” if it contained at least two cells distributed across a given cell-type pair {*s*1, *s*2}. A cumulative sharing metric was calculated by summing the fractional representation 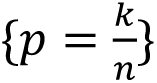 of all shared clones, where *k* is the number of clonal cells in a particular cell type and > is the total cell number belonging to this specific clone across all cell types. To determine statistical significance, 10,000 permutations were performed by randomly shuffling cell-type assignments while preserving the original cluster size distribution. The lineage coupling z-score was then derived by comparing the observed metric against this null distribution. Lineage coupling correlations were further calculated as the Pearson correlation coefficients between the z-score profiles of each cell-type pair.

### Spatial distance and transcriptomics distance between clone sister cells

To assess the spatial and transcriptomic cohesion of cell clones, we performed a distance-based analysis comparing clone-specific pairs against random cellular backgrounds. The spatial dispersion of each clone was quantified by calculating the Euclidean distance between the physical coordinates (x, y) of all cell pairs within the same clone. To establish a null distribution, a random background was generated by calculating the Euclidean distances between an equivalent number of randomly sampled cell pairs from the entire clone dataset. Transcriptomic similarity within clones was evaluated by computing the Euclidean distance between clone pairs in the two-dimensional UMAP embedding (UMAP_1 and UMAP_2). A corresponding random background was established by calculating the distances between randomly paired cells within the same UMAP space.

### Spatial clonal orientation analysis

To quantify the spatial orientation of clonal expansion relative to anatomical structures, we calculated the alignment angles between clonal sister pairs and a defined tissue horizontal axis. Tissue horizontal axis was represented as a series of smoothed and ordered coordinates of specific regional curvilinear geometry based on L2 subregion annotation. For any given location in the structure, the local orientation was defined as the tangent vector of the tissue axis curve (*v_a_*). Specifically, for a cell pair link-line (*v_b_*), we identified the nearest point Pab on the curve of its midpoint. The angle θ between the clonal sister the local axis was calculated using the dot product formula: 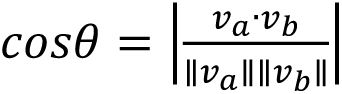

## Data availability

All sequencing datasets generated in this study are deposited under BioProject accession number PRJNA1436734. Processed spatial transcriptomics and lineage data are available upon email request.

## Code availability

All source code is available at https://github.com/gpenglab/Steller.

## Acknowledgements

We thank Min Liu, Menglong Zhang and Yuntong Wang from BMKMANU company for help with modifying spatial chip with CS probes and for critical discussion about data analysis; Rongyang Li from Weike Pei’s lab at Westlake University for help with CloneBC plasmid and lentivirus preparation as well as 4T1 tumor construction; GIBH Analytical Instrumentation Core for their support with FACS; and Feng Yang from Bin Zhang’s lab at Westlake University for guidance on in utero injections. This study was supported by grants from the National Key R&D Program of China (2024YFC3405600, 2022YFA1105700), National Natural Science Foundation of China (32270854, 32450525, 82270123), Guangdong Major Project of Basic Research (2026B0303000013), Guangdong Basic and Applied Basic Research Foundation (2024B1515040020), Science and Technology Planning Project of Guangdong Province (2023B1212060050, 2023B1212120009), Basic Research Project of Guangzhou Institutes of Biomedicine and Health, Chinese Academy of Sciences (GIBHBRP23-01, GIBHBRP24-01), Major Project of Guangzhou National Laboratory (GZNL2023A03005), Zhejiang Provincial Natural Science Foundation of China (LR25C070001), Westlake Genome Editing Cutting-Edge Exploration Program (21200000A992510/002), Pioneer and Leading Goose Key R&D Program of Zhejiang Province (2024SSYS0033, 2024SSYS0034), Guangdong Provincial Key Laboratory of Stem Cell and Regenerative Medicine (KLRB202303).

## Author Contributions

Z.L. led experimental work, generated and analyzed data, and prepared figures, supervised by G.P. and W.P.; Experiments were collaborated with C.W., C.G., C.C., Y.L. and Q.X.; Z.L. C.W. and G.P. wrote the manuscript, with help from all authors.

---

**Figure S1.**
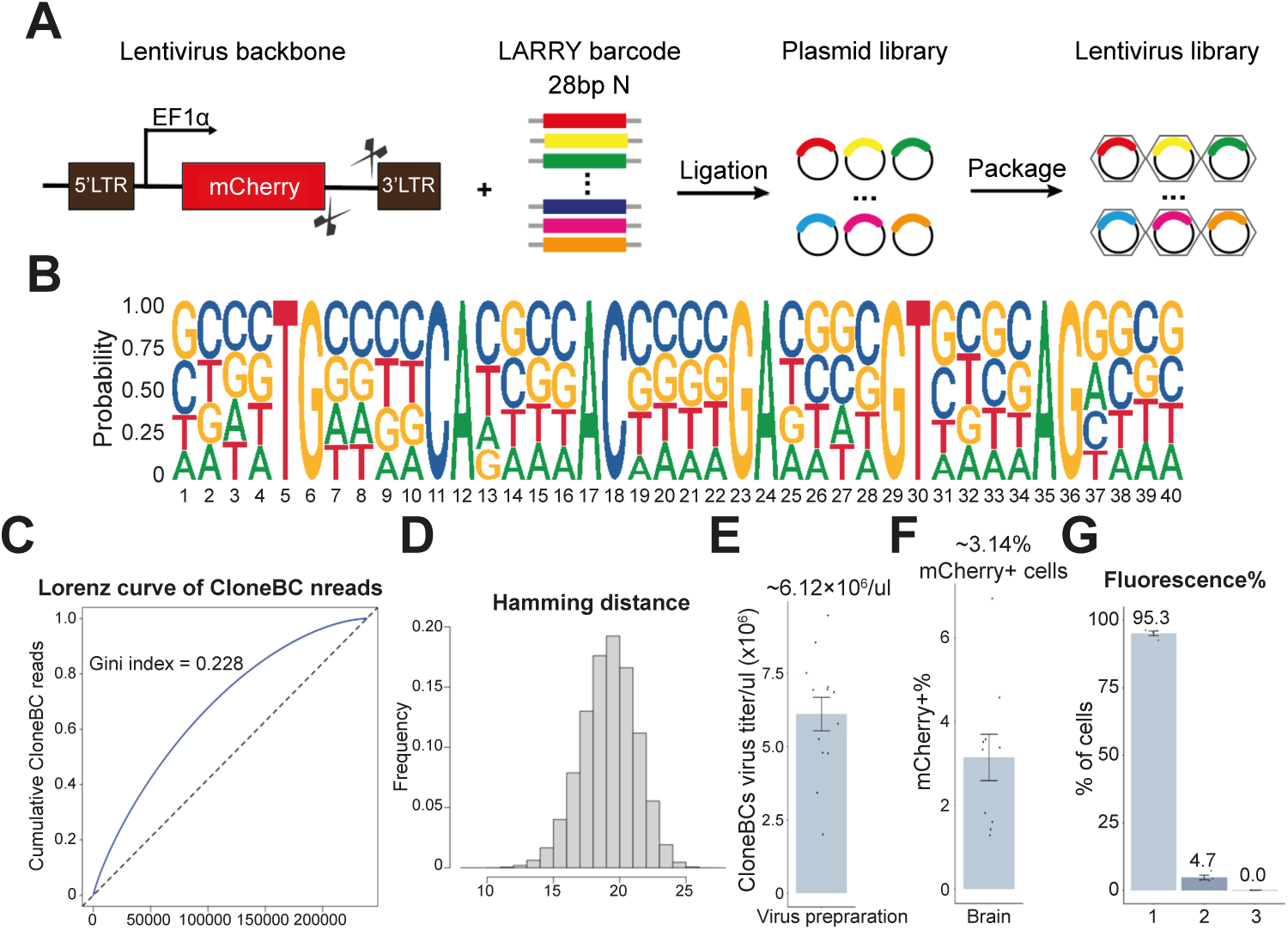
Construction of CloneBC plasmid library and lentivirus for *in utero* injection. **(A)** Experimental workflow for the construction of CloneBC plasmid and lentiviral libraries. **(B)** Sequence logo depicting the nucleotide composition and position-specific probability of the CloneBC library. **(C)** Cumulative distribution of CloneBC read abundances within the plasmid library whitelist. The diagonal dashed line represents a theoretical perfectly uniform distribution. **(D)** Distribution of pairwise Hamming distances among CloneBC sequences within the plasmid library, indicating high sequence diversity and error-correction potential. **(E)** Bar plot showing lentiviral titers across independent preparations (mean = 6.12 x 10^9 TU/µL, n = 21). Mean ± SEM. **(F)** Flow cytometric quantification of mCherry+ cell percentages across replicates (mean = 3.14%, n = 11). Mean ± SEM. **(G)** Proportion of cells labeled with single-color versus dual-color fluorescence at E14.5–E15.5 following E11.5 microinjection (n = 4). Mean ± SEM.

**Figure S2.**
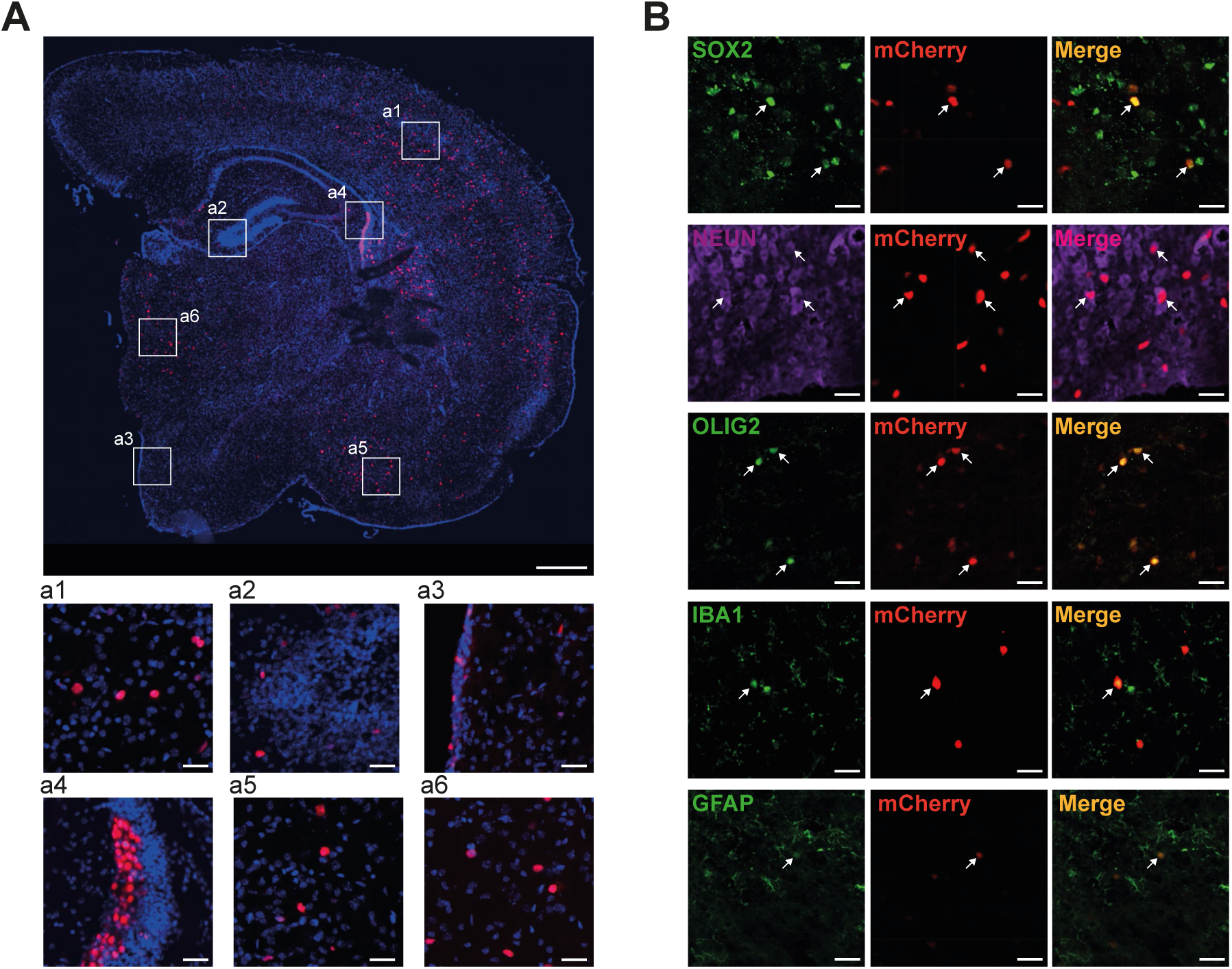
Cell fate identification and distribution of CloneBC lentivirus traced postnatal progenies. **(A)** Representative coronal section of a P10 mouse brain following in utero CloneBC lentiviral injection at E12.5, showing widespread distribution of mCherry-labeled clonal descendants. High-magnification views of the boxed regions a1–6 (bottom) illustrate labeled cells in the cortex (a1), hippocampus (a2), hypothalamus (a3), lateral ventricular zone (LVZ; a4), subcortex (a5), and thalamus (a6). **(B)** Immunofluorescence characterization of clonal cell identities in P10 coronal sections. Co-staining of mCherry with lineage-specific markers: SOX2 (progenitors), NEUN (neurons), OLIG2 (oligodendrocytes), IBA1 (microglia), and GFAP (astrocytes). Arrowheads indicate representative cells co-expressing mCherry and the respective markers. Scale bars: 500 μm in (A, upper), 30 μm in (A, lower), and 10 μm in (B).

**Figure S3.**
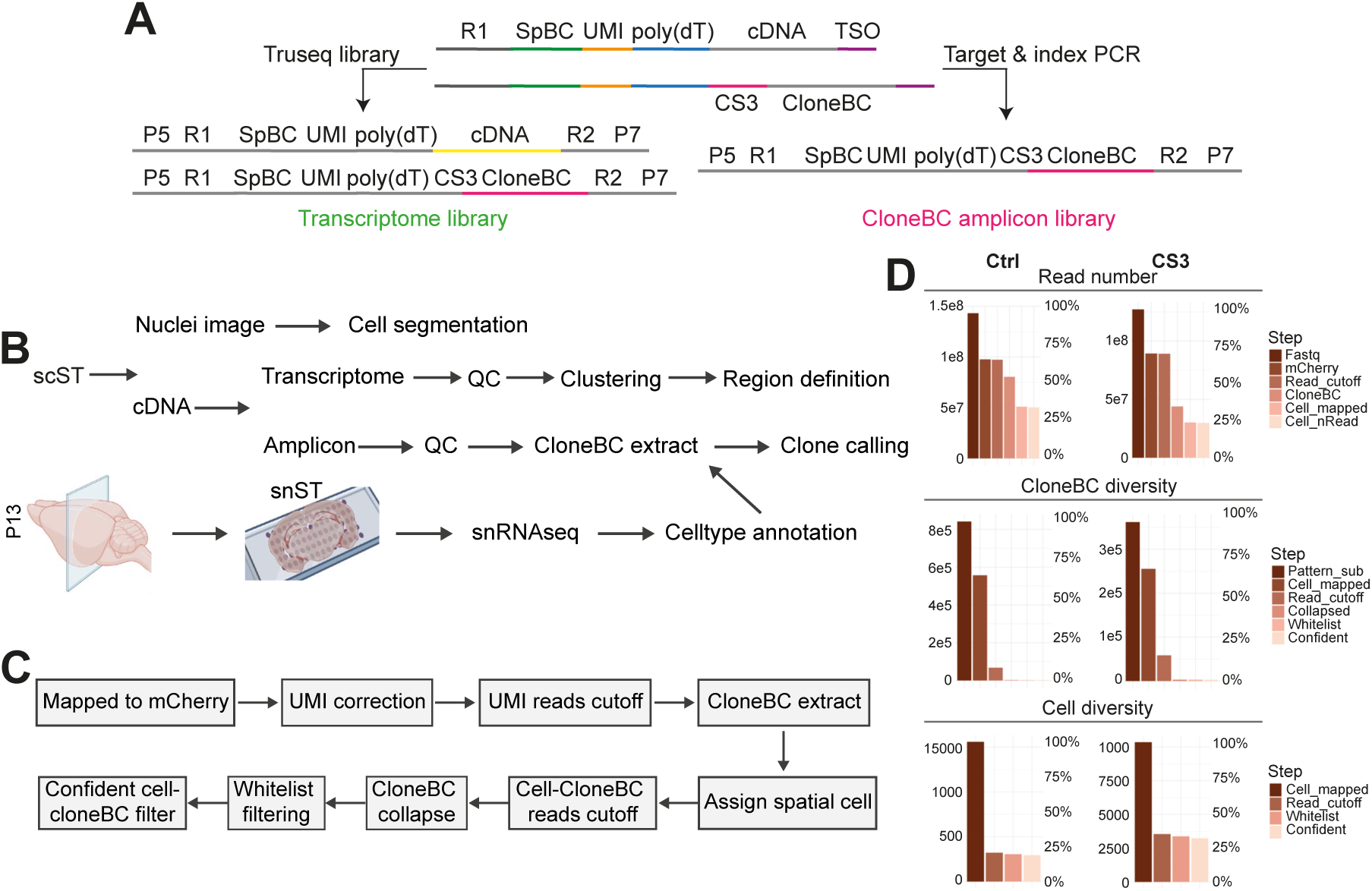
Transcriptome and CloneBC library generation, analytical pipeline and quality control. **(A)** Schematic workflows for spatial library construction. Left: whole-transcriptome library preparation via the TruSeq protocol; Right: target CloneBC enrichment via site-specific PCR amplification. **(B)** Computational pipeline for the integration of multimodal spatial datasets, combining whole-transcriptome and targeted CloneBC amplicon data, see Methods for details. **(C)** Schematic of the bioinformatics workflow for processing CloneBC amplicon sequences, including barcode extraction, filtering, and assignment, see Methods for details. **(D)** Step-wise retention analysis of total reads, cell diversity (number of unique cells), and unique CloneBCs throughout the pre-processing pipeline for ctrl and CS3 libraries.

**Figure S4.**
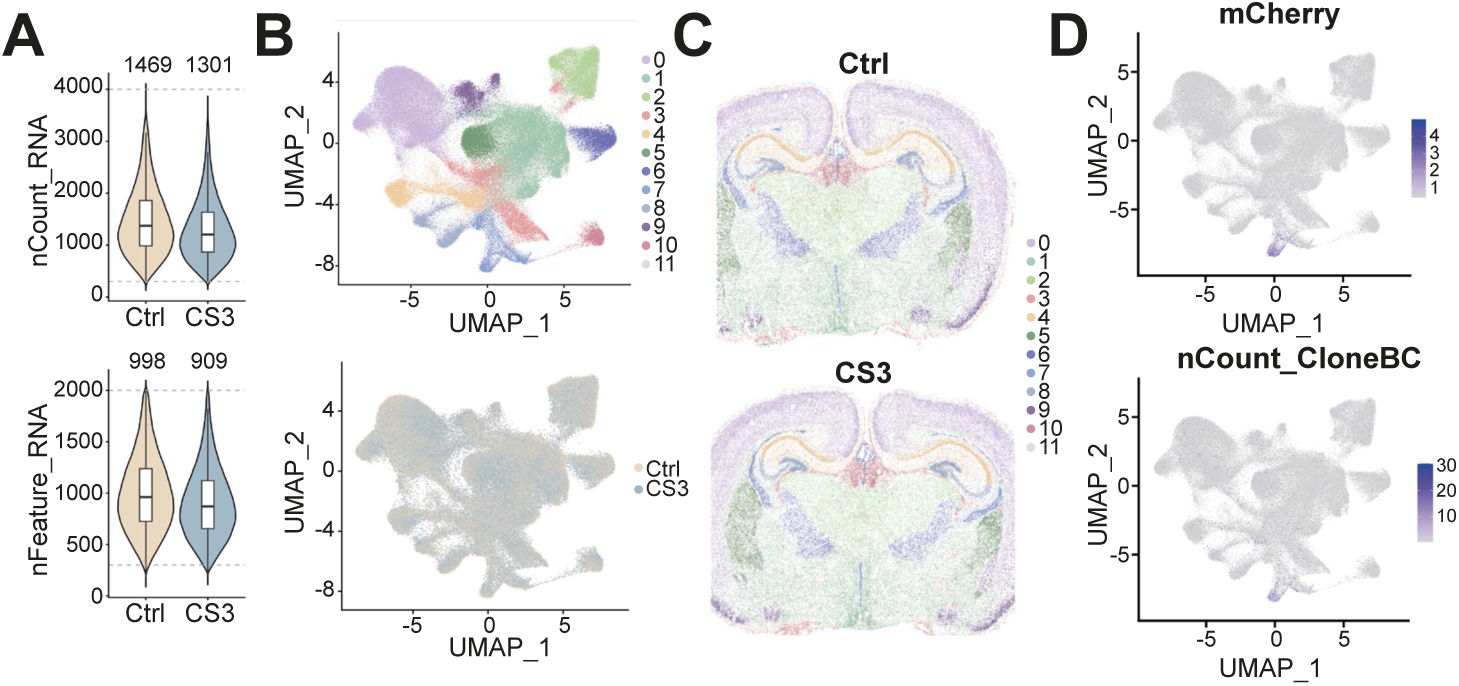
Transcriptomic quality control. **(A)** Violin plots of unique molecular identifier (UMI) counts (nCount) and detected genes (nFeature) per cell for ctrl and CS3 sections. Mean values for each metric are indicated. **(B)** UMAP projections of integrated cells, color-coded by unsupervised clusters (top) and sample groups (bottom), demonstrating high transcriptomic comparability between ctrl and CS3 datasets. **(C)** Spatial distribution of identified cell clusters across the two tissue sections, reflecting anatomical consistency. **(D)** Feature plots comparing the spatial counts of mCherry transcripts (from whole-transcriptome equencing) and CloneBC tags (from targeted amplicon sequencing) in the same tissue sections.

**Figure S5.**
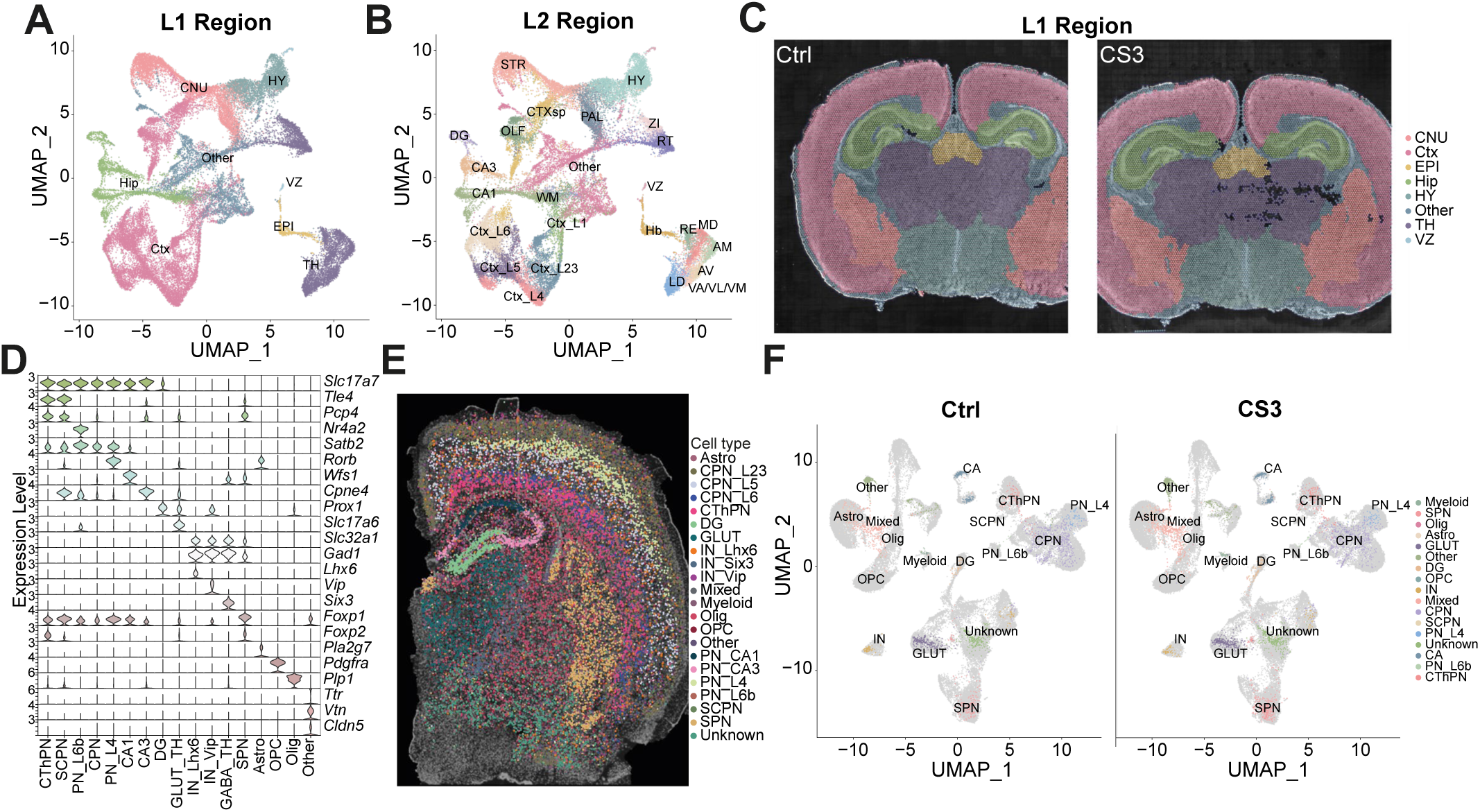
Anatomical region delineation and cell-type distribution. (A-C) Multi-level anatomical annotation of the mouse brain visualized via UMAP (A, B) and spatial maps (C). Brain regions are delineated at two resolutions: Level 1 (L1) represents broad anatomical domains (a, c), while Level 2 (L2) provides fine-scale sub-structural resolution (B). Cortex (Ctx); Cerebral nuclei (CNU); Hippocampus (Hip); Thalamus (TH); Hypothalamus (HY); Epithalamus (EPI); Ventricular zone (VZ). **(D)** Expression of canonical marker genes across identified cell types in the integrated dataset. **(E)** Spatial distribution of identified cell types across representative snST sections. **(F)** Split-UMAP projections of major cell types categorized by sample groups, demonstrating consistency in cell-type recovery. Astro: Astrocyte; CPN_L23: Layer 2/3 callosal pyramidal neuron; CThPN: corticothalamic projection neuron; DG: dentate gyrus granule cell; IN: interneuron; Olig: oligodendrocyte; OPC: oligodendrocyte precursor cell; PN: pyramidal neuron; SCPN: subcerebral projection neuron; SPN: spiny projection neuron; GLUT: glutamatergic neuron.

**Figure S6.**
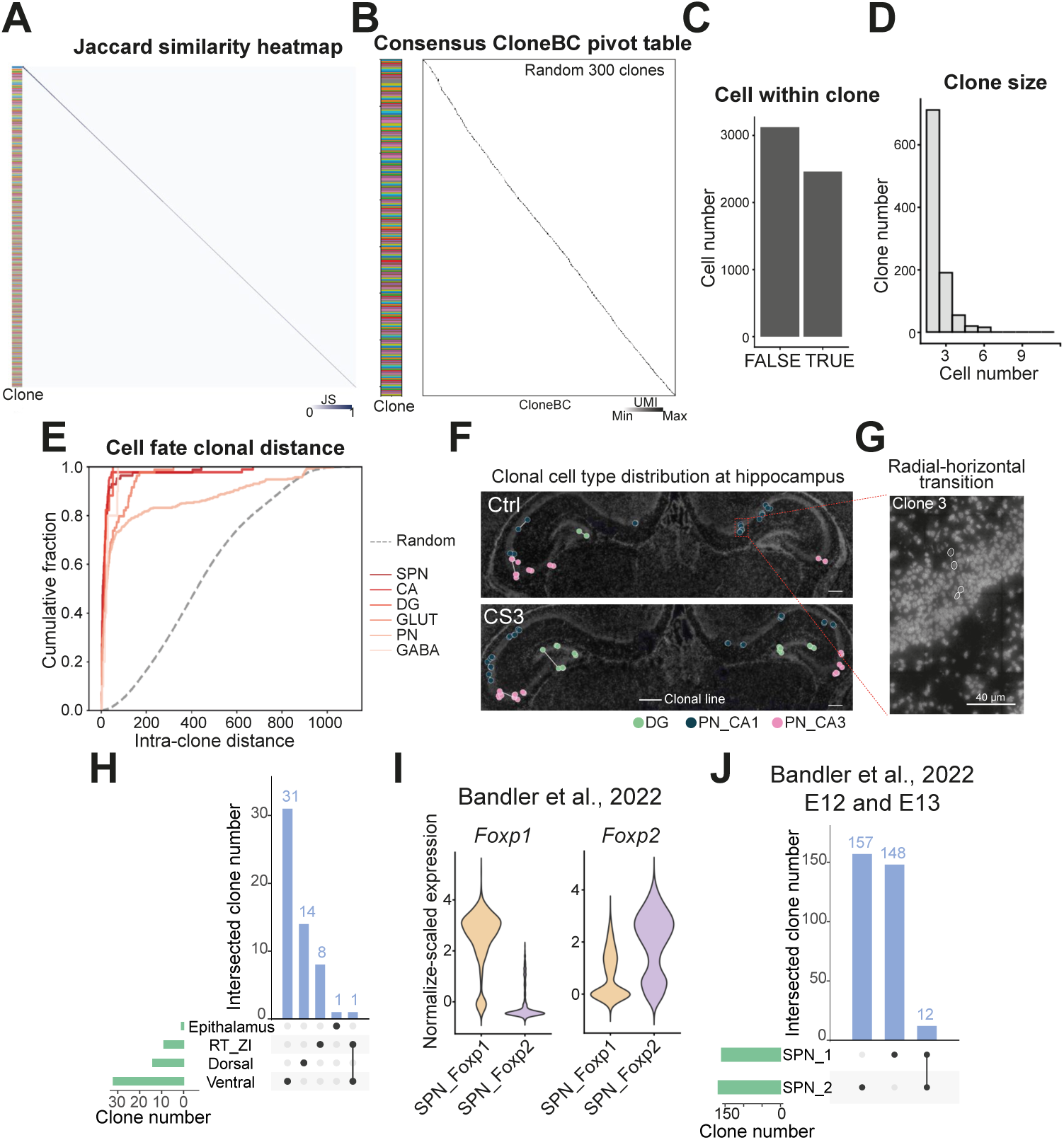
High-fidelity clonal identification and spatial lineage fate. **(A)** Heatmap of Jaccard similarity indices for CloneBC combinations within individual spatial cells, demonstrating the uniqueness of barcode signatures. **(B)** Pivot-table heatmap of the cell-by-CloneBC UMI count matrix. Rows and columns are organized by assigned clonal identities (indicated by row-side color bars). The high-intensity diagonal blocks represent robust UMI counts per cell-CloneBC pair, highlighting the specificity and stringency of clonal assignment. **(C)** Bar plot showing the distribution of clones based on the number of detected sister cells per clone. **(D)** Histogram illustrating the distribution of clonal sizes (cell count per clone) across the dataset. **(E)** Cumulative distribution functions (CDFs) comparing the spatial proximity of clonal sister pairs within major lineages against randomly sampled cell pairs, indicating significant spatial clustering of related cells. SPN: spiny projection neuron; CA: CA1 and CA3 pyramidal neuron; DG: DG granule neuron; GLUT: Glutamatergic neuron; PN: cortical pyramidal neuron; GABA: GABAergic neuron. **(F)** Representative spatial maps illustrate the organization of neuronal clones across the CA1, CA3, and DG regions in two independent tissue sections. Cells are color-coded by their respective cell-type identities, and sister cells are bridged by lines of corresponding colors. **(G)** Cell outline white mask of clone 3 overlaid with dapi staining image to show the radial-to-horizontal transition migration pattern at P4 mouse CA1 region. **(H)** UpSet plot showing the intersection of unique clones across discrete regions of the TH and Hb. **(I-J)** External validation of SPN molecular and clonal features using the dataset from Bandler et al. (2022). Violin plots (I) show *Foxp1* and *Foxp2* expression levels across SPN subtypes, and the UpSet plot (J) depicts the clonal intersection between these subtypes from dataset labeled at E12 and E13. Scale bars: 200 μm in (F) and 40 μm in (G).

